# Multimodal Object Representations Rely on Integrative Coding

**DOI:** 10.1101/2022.08.31.504599

**Authors:** Aedan Y. Li, Natalia Ladyka-Wojcik, Heba Qazilbash, Ali Golestani, Dirk B. Walther, Chris B. Martin, Morgan D. Barense

## Abstract

Combining information from multiple senses is essential to object recognition. Yet how the mind combines sensory input into coherent multimodal representations – the *multimodal binding problem* – remains poorly understood. Here, we applied multi-echo fMRI across a four-day paradigm, in which participants learned 3-dimensional multimodal object representations created from well-characterized visual shape and sound features. Our novel paradigm decoupled the learned multimodal object representations from their baseline unimodal shape and sound features, thus tracking the emergence of multimodal concepts as they were learned by healthy adults. Critically, the representation for the whole object was different from the combined representation of its individual parts, with evidence of an integrative object code in anterior temporal lobe structures. Intriguingly, the perirhinal cortex – an anterior temporal lobe structure – was by default biased towards visual shape, but this initial shape bias was attenuated with learning. Pattern similarity analyses suggest that after learning the perirhinal cortex orthogonalized combinations of visual shape and sound features, transforming overlapping feature input into distinct multimodal object representations. These results provide evidence of integrative coding in the anterior temporal lobes that is distinct from the distributed sensory features, advancing the age-old question of how the mind constructs multimodal objects from their component features.

---

The world is a great blooming, buzzing confusion^1^ of the senses. Our ability to understand “what is out there” depends on combining sensory features to form *multimodal object concepts.* A child, for example, might form the concept “frog” by learning that the visual appearance of a four-legged creature goes with the sound of its croaking. Consequently, this child has also learned that frogs do not produce barking sounds, as the child has created a unique object association from specific shape and sound features. Forming coherent object representations is thus essential for human experience, allowing the child to behave adaptively under changing environments. Yet how is it possible for the child to know that the sound of croaking is associated with the visual shape of a frog, even when she might be looking at a dog? How does the human mind form meaningful concepts from the vast amount of feature information that bombards the senses, allowing us to interpret our external world?

Known as the *multimodal binding problem,* an unresolved question in the cognitive sciences has asked how the mind combines sensory features into coherent multimodal object representations. One theoretical view predicts that multimodal objects are built from component unimodal features represented across distributed sensory regions.^2^ Under this view, when a child thinks about “frog”, the visual cortex represents the appearance of the frog whereas the auditory cortex represents the croaking sound. Alternatively, other theoretical views predict that objects are not only built from sensory features, but that there is also an explicit integrative code distinct from the features (i.e., the whole is different than the sum of its parts).^3,4,5,6,7^ These latter views propose that the anterior temporal lobes act as a polymodal “hub” that combine separate features into integrated wholes.^3,5,8,9^ Thus, the key theoretical challenge central to resolving the multimodal binding problem is understanding the format of the object representation. Are multimodal objects entirely built from features distributed across sensory regions, or is there also an explicit integrative code distinct from component features in the anterior temporal lobes? Furthermore, the existing literature has predominantly studied the neural representation of well-established object concepts from the visual domain alone,^2-19^ even though human experience is fundamentally multimodal.

Here, we leveraged multi-echo fMRI^20^ across a novel four-day task in which participants learned to associate unimodal visual shape and sound features into 3D multimodal object representations. First, we characterized shape^21^ and sound features in a separate validation experiment, ensuring that the features were well-matched in subjective similarity (*Figure 1*). On the learning task, participants independently explored the 3D-printed shapes and heard novel experimenter-constructed sounds. The participants then learned specific shape-sound associations (congruent objects), while other shape-sound associations were not learned (incongruent objects). Critically, these learned and non-learned objects were constructed from the same set of shape and sound features, ensuring that the participant familiarity with individual features were tightly controlled. With this design, our four-day learning task explored whether the learned multimodal object representations may be transformed from their original unimodal features (i.e., whether the conjunction of multimodal features is different than the sum of their individual parts).

**Figure 1.**
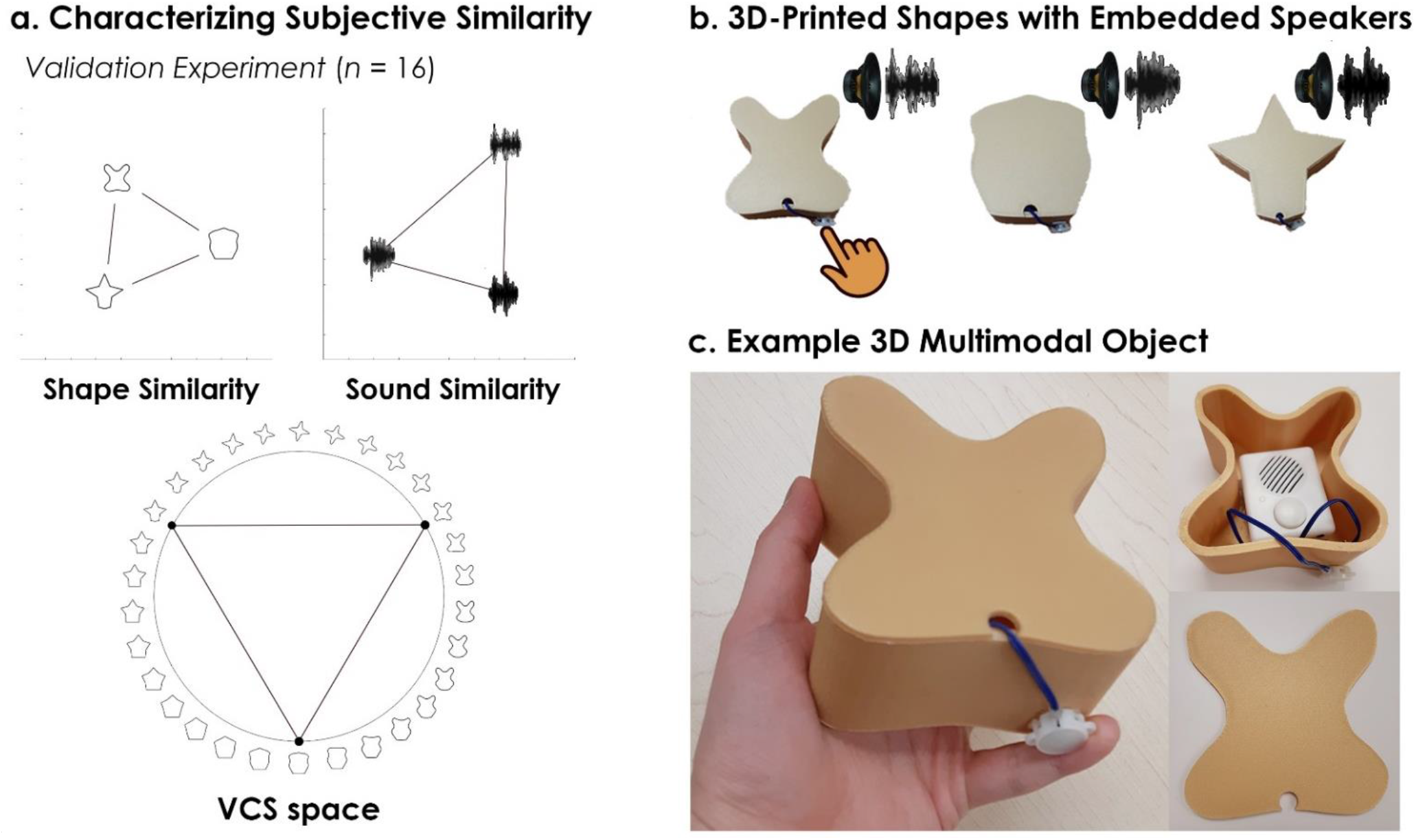
D-printed objects. An independent validation experiment ensured that the similarity of shapes and sounds were well-matched before the learning task. (*a*) Three shapes were sampled from the *Validated Circular Shape (VCS) Space* (shown as black points on VCS space),^21^ a stimulus space whereby angular distance corresponds to subjective shape similarity. Three sounds were sampled from a validated set of five experimenter-created sounds. As shown by the triangular representational geometries above, we confirmed that participants rated the shapes to be as similar to other shapes as sounds were to other sounds (visualized using multidimensional scaling^22^; also see *Supplemental Material Figure S1*). This independent validation experiment ensured that we could characterize the change in similarity structure following multimodal learning, because we knew the baseline similarity structure (i.e., two triangular representational geometries). (*b*) The shapes were 3D-printed with a hollow space and embedded with a button-activated speaker. (*c*) Participants could physically explore and palpate the 3D multimodal objects. Critically, we manipulated whether the button-activated speaker was operational across learning days (see Methods/Figure 2).

We found that multimodal object concepts were represented as distributed sensory-specific features along the visual and auditory processing pathways, as well as explicit integrative combinations of those features in the anterior temporal lobes. Intriguingly, the perirhinal cortex – an anterior temporal lobe structure – was biased towards the visual modality before multimodal learning, with greater activity towards shape over sound features. Pattern similarity analyses revealed that the shape representations in perirhinal cortex were initially unaffected by sound, providing evidence of a default visual shape bias. However, this initial visually-biased code came to be transformed with experience, as multimodal object representations (measured after multimodal learning) differed from component feature representations (measured before multimodal learning).

In summary, these results provide new insights towards an age-old problem: we show that the mind constructs multimodal object concepts through an explicit integrative code in the anterior temporal lobes, which is distinct from the component object features that are distributed across sensory cortices.

## Results

### Multimodal Object Learning

We first characterized the similarity structure of unimodal features, asking an independent group of 16 participants to rate the similarity of shapes^21^ and sounds (*Figure 1a,* see *Supplemental Material Figure S1*). This procedure ensured that the subjective similarity of features was well-matched before the primary learning task (*Figure 2*). The visual shape features were then 3D-printed and embedded with button-activated programmable speakers (*Figure 1b*). During the task, each participant learned a set of shape-sound associations (congruent objects). Before multimodal learning, participants explored the 3D-printed shapes and sounds separately (*Day 1:* the button-activated speaker was not operational, although participants pressed the button as they explored the shapes). After multimodal learning, participants explored specific shape-sound associations, with pairings counter-balanced across observers (*Day 3:* by pressing the button on each 3D-printed shape to play the associated sound).

**Figure 2.**
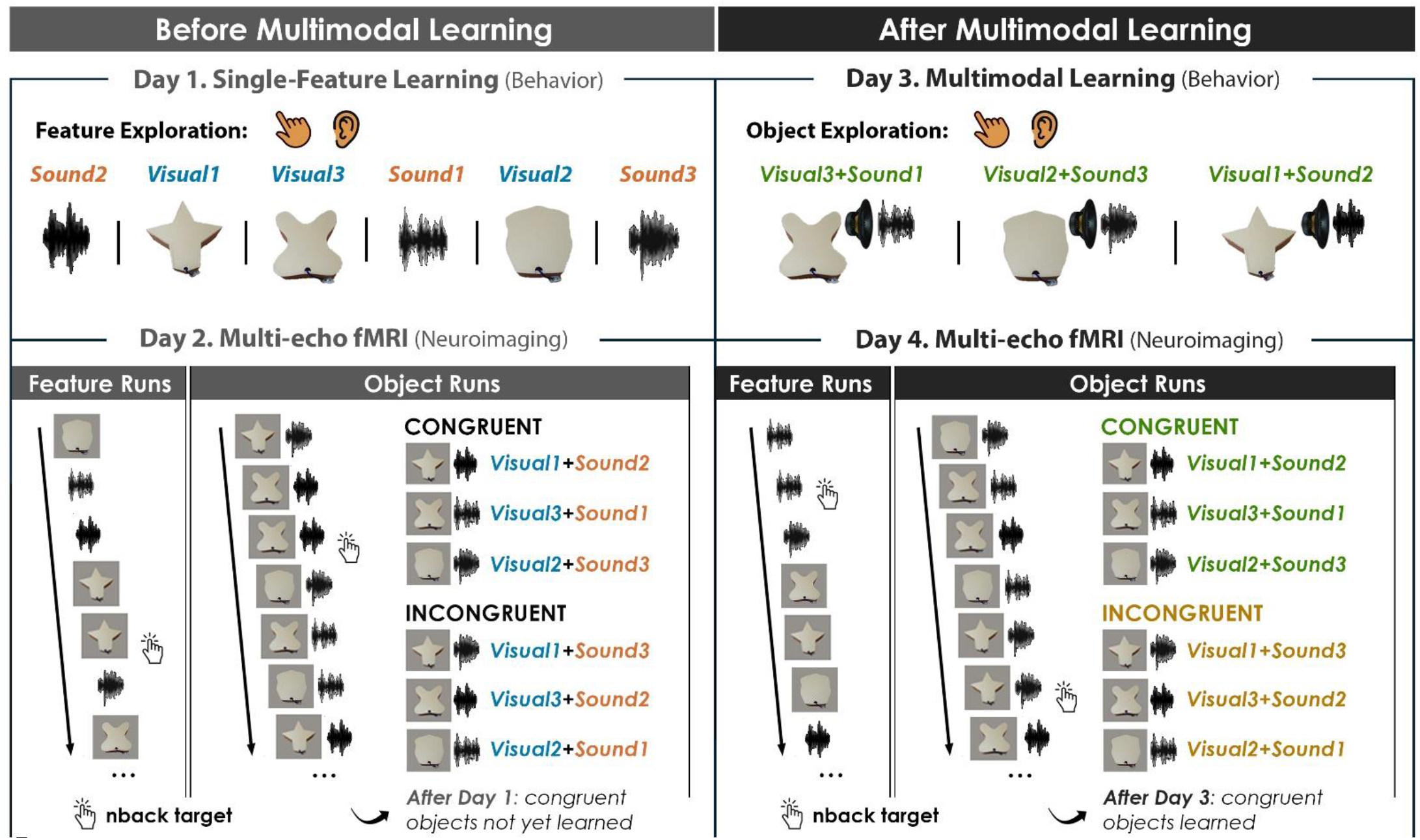
Four-day multimodal object learning task. On **Day 1** (behavior), participants explored 3D-printed shapes while the button-activated speakers were not operational and heard sounds through a headset. During a separate task, participants rated the similarity of the visual shapes and sound features. On **Day 2** (neuroimaging), participants completed 10 feature runs in which shape and sound unimodal features were experienced separately and 5 object runs in which shape and sound were experienced simultaneously. As participants have not yet learned the congruent shape-sound pairings, the Day 2 neuroimaging session serves as a within-subject neural baseline for how the unimodal features were represented before multimodal learning. On **Day 3** (behavior), participants again explored the shape and sound features. Participants now learned to associate specific visual and sound features together as a multimodal object by pressing the button to play an embedded speaker, forming congruent object representations (i.e., multimodal learning). Shape-sound associations were counterbalanced across participants, and we again collected similarity ratings between the shapes and sounds on a separate task. On **Day 4** (neuroimaging), participants completed the same task as on Day 2. Across four days, we characterized the neural and behavioral changes that occurred before and after shapes and sounds were paired together to form multimodal object representations. As the baseline similarity structure of the shape and sound features were *a priori* defined (see *Figure 1*) and measured on the first day of learning (see *Supplemental Figure S1*), changes to the within-subject similarity structure provide insight into whether the object representations (acquired after multimodal learning) differed from component feature representations (acquired before multimodal learning).

On neuroimaging days (*Day 2* and *Day 4*), we recorded neural responses to features presented separately and to features presented simultaneously using multi-echo fMRI (*Figure 2*). During feature runs, participants either viewed images of 3D-printed shapes (*Visual*) or heard sounds (*Sound*). During object runs, participants experienced either the shape-sound associations learned on Day 3 (*Congruent*) or shape-sound associations that had not been learned on Day 3 (*Incongruent*). All participants could recognize their specific associations at the end of Day 3, confirming that the congruent shape-sound objects were successfully learned (performance = 100% for all participants).

### Unimodal Shape and Sound Representations are Distributed

#### Whole-brain Univariate Analysis

In the first set of neuroimaging analyses, we examined whether distributed brain regions were involved in representing unimodal shapes and sounds. During feature runs (shapes and sounds presented separately), we observed robust bilateral modality-specific activity across the neocortex (*Figure 3a-c*). The ventral visual stream extending into the perirhinal cortex activated more strongly to visual compared to sound features (*Figure 3a*). The auditory processing stream, from the primary auditory cortex extending into the temporal pole along the superior temporal sulcus, activated more strongly to sound compared to visual features (*Figure 3b*). These results provide an important baseline, replicating the known representational divisions across the neocortex.^23,24,25^

**Figure 3.**
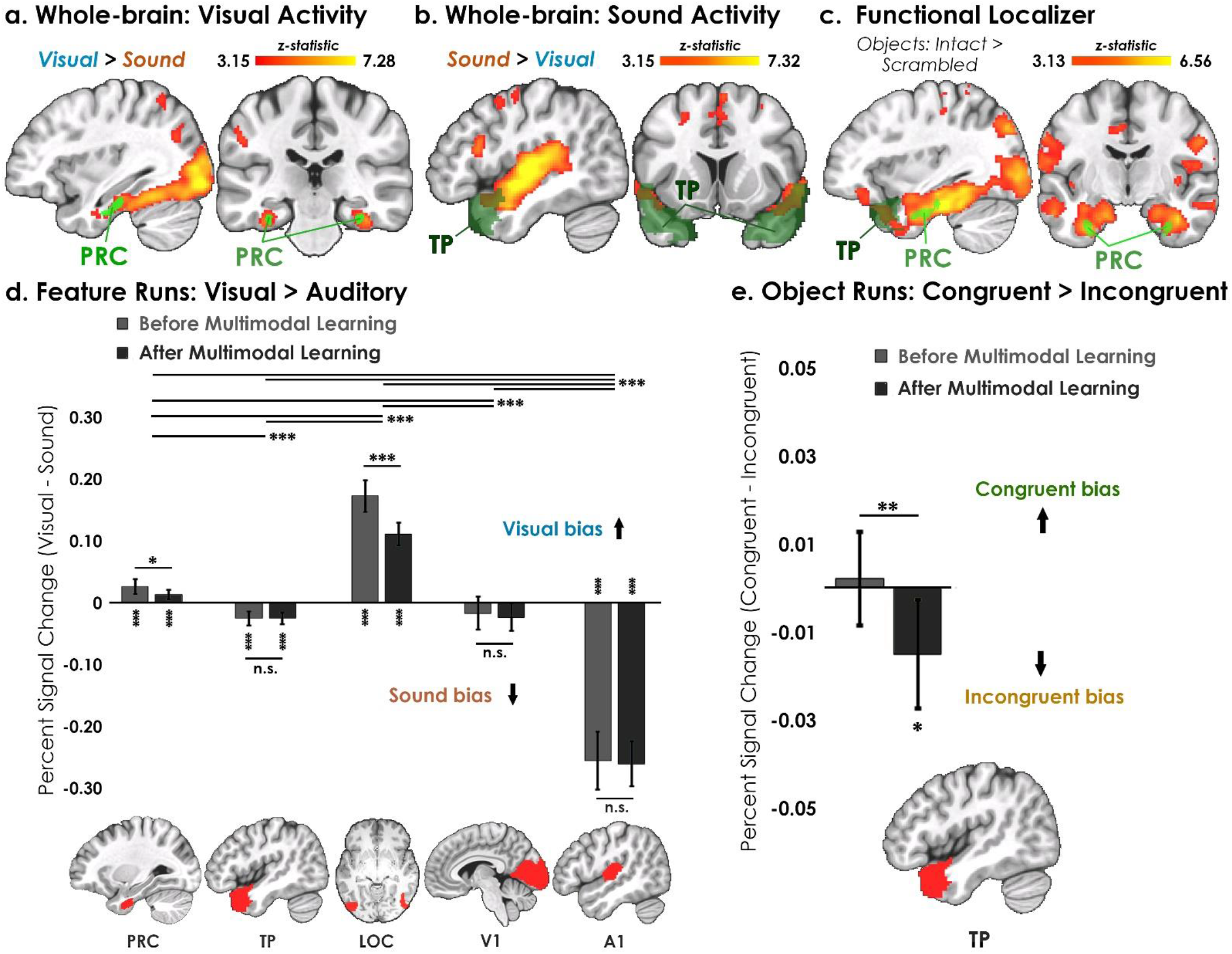
Univariate analyses superimposed on MNI-152 standard space. All contrasts were thresholded at voxelwise *p* = 0.001 and cluster-corrected at *p* = 0.05 (random-effects, FSL FLAME; 6-mm spatial smoothing). Five ROIs were *a priori* selected based on existing theory: temporal pole – TP, perirhinal cortex – PRC, lateral occipital complex – LOC, primary visual cortex – V1, and primary auditory cortex – A1. (*a-c*) Collapsed across learning days, robust modality-specific activity was observed across the neocortex. (*d*) Consistent with the whole-brain results, LOC was biased towards visual features whereas A1 was biased towards sound features. Intriguingly, we found learning-related shifts in activation in PRC and LOC, with the magnitude of visual bias decreasing after multimodal learning. Critically, whereas voxels in the (*d*) TP mask were biased towards sound information, (*e*) voxels in the same TP mask also reflected an experience-dependent reduction in activity to learned shape-sound associations. As participants experienced the same shape and sound features before and after multimodal learning, these results provide neural evidence of an explicit multimodal integrative code in the temporal pole. Notably, there were no experience-dependent univariate activity differences in any other examined region during object runs. * denotes *p* < 0.05, ** denotes *p* < 0.01, *** denotes *p* < 0.001. Horizontal lines at top of figure denotes significant differences across brain regions (e.g., LOC is more biased towards visual shape than any other region), whereas horizontal lines within brain regions reflect an effect of learning day (e.g., reduction in visual bias after multimodal learning in PRC).

Complementing these results, a standard functional localizer revealed greater activation for intact objects compared to phase scrambled objects along the ventral visual stream, extending into medial temporal and anterior temporal lobe regions (*Figure 3c*). Supporting previous theory predicting perirhinal cortex involvement in visual object representation at short delays,^4,5,6^ these results indicate perirhinal cortex activity was by default biased towards visual information in the feature runs (i.e., towards complex visual shape configurations; *Figure 3a*) and towards intact objects rather than scrambled objects in the functional localizer (*Figure 3c*).

Next, we determined how neural responses changed following multimodal learning. We compared the associations learned by participants to associations that were not learned (Object runs: *Congruent > Incongruent* contrast). No regions survived thresholding for the *Congruent > Incongruent* contrast, so we examined univariate activity within five regions thought to be important for representing features and their integration: temporal pole, perirhinal cortex, lateral occipital complex (LOC), primary visual cortex (V1), and primary auditory cortex (A1).^3,5^

#### Region-of-Interest Univariate Analysis

We observed an interaction between learning day (before or after multimodal learning) and modality (visual or sound feature) in perirhinal cortex (*F*_1,48_ = 4.36, *p* = 0.027, *η*^2^ = 0.098; *Figure 3d*) and LOC (*F*_1,45_ = 25.89, *p* < 0.001, *η*^2^ = 0.37). These regions activated more strongly to visual information before multimodal learning compared to after multimodal learning, indicative of a visual bias that was attenuated with experience. Across days, perirhinal cortex (*t*_67_ = 5.53, *p* < 0.001, *Cohen’s d* = 0.67) and LOC (*t*_63_ = 16.02, *p* < 0.001, *Cohen’s d* = 2.00) were biased towards visual information, whereas temporal pole (*t*_67_ = 6.73, *p* < 0.001, *Cohen’s d* = 0.82) and A1 (*t*_67_ = 17.09, *p* < 0.001, *Cohen’s d* = 2.07) were biased towards sound information. Interestingly, we found a small overall bias towards sound in V1^26^ (*t*_67_ = 2.26, *p* = 0.027, *Cohen’s d* = 0.20).

### Multimodal Objects Rely on Integrative Coding in the Anterior Temporal Lobes

In the final univariate analysis, we examined the *Congruent > Incongruent* contrast in object runs (*Figure 2b, d*). A significant interaction was found between learning day (before or after multimodal learning) and congruency (congruent or incongruent features) in the temporal pole (*F*_1,48_ = 7.63, *p* = 0.0081, *η*^2^ = 0.14). Critically, there was no difference in activity between congruent and incongruent objects before multimodal learning (*t*_33_ = 0.37, *p* = 0.72), but there was more activation to incongruent compared to congruent objects after multimodal learning (*t*_33_ = 2.42, *p* = 0.021, *Cohen’s d* = 0.42).

As the shape-sound features experienced by participants were the same before and after multimodal learning (*Figure 2*), these neural changes imply the formation of an explicit integrative code in the temporal pole. That is, the temporal pole differentiated congruent and incongruent objects built from the same features, indicative of a multimodal object code distinct from the sum of component feature parts. By contrast, we did not observe a difference between congruent and incongruent objects in perirhinal cortex, LOC, V1, or A1 (*F*_1,45~48_ = 0.088~2.34, *p* = 0.13~0.77).

The previous univariate analyses provide evidence of feature coding distributed across visual and auditory processing streams (*Figure 3a-d*), as well as multimodal integrative coding within the anterior temporal lobes (*Figure 3e*). However, the univariate evidence of explicit integrative coding in the anterior temporal lobes may be driven by a novelty signal *per se* rather than by an integrative code specific to individual object concepts. To better understand the representational format of the multimodal objects, we analyzed the within-subject change in similarity structure *before* and *after* multimodal learning. In other words, we determined whether initial feature representations (before multimodal learning) changed after they were combined into object representations (after multimodal learning), providing evidence of experience-dependent transformations specific to each object that may occur with learning.

We first conducted the pattern similarity analyses in behavior, examining how participants experienced the shapes and sounds across learning days. We then conducted a series of neuroimaging pattern similarity analyses across feature-and object-runs before and after multimodal learning. One prediction may be that unimodal feature representations are not affected when they are paired together to form multimodal object representations (i.e., object concepts are entirely built from component features). Alternatively, unimodal feature representations may change after they are paired to form multimodal objects, such that multimodal learning transforms the original feature representations (i.e., acquired object concepts are abstracted away from the original component features).

#### Behavioral Pattern Similarity Results

There was a robust learning-related change in how participants experienced the similarity of shape and sound features (*Supplemental Material. Figure S1, S2*). In behavior, there was a main effect of learning day (before or after multimodal learning: *F*_1,51_ = 24.45, *p* < 0.001, *η^2^* = 0.32), a main effect of congruency (congruent or incongruent: *F*_1,51_ = 6.93, *p* = 0.011, *η*^2^ = 0.12), and an interaction between learning day and congruency (*F*_1,51_ = 15.33, *p* < 0.001, *η*^2^ = 0.23). Before multimodal learning, there was no difference in similarity between congruent and incongruent shape-sound features (*t*_17_ = 0.78, p = 0.44). Furthermore, subjective similarity was approximately matched for both shape and sound (i.e., two triangular representational geometries, replicating the independent validation experiment, *Supplemental Material. Figure S1a, S2a*). After multimodal learning (*Supplemental Material. Figure S1b, S2b*), participants rated shapes and sounds associated with congruent objects to be more similar than shapes and sounds associated with incongruent objects (*t*_17_ = 5.10, *p* < 0.001, *Cohen’s d* = 1.28), an effect observed in 17/18 participants. We confirmed this experience-dependent change in similarity structure in a separate behavioral experiment with a larger sample size (observed in 38/44 participants; learning day × congruency interaction: F_1,129_ = 13.74, p < 0.001; *η*^2^ = 0.096; *Supplemental Material. Figure S2*).

As the physical features themselves had not changed across the learning task, this within-subject change in representational geometry provides evidence that multimodal learning changed how participants subjectively experienced objects constructed from combinations of the same features.

#### Neural Pattern Similarity Results

##### Explicit integrative coding: Congruent features differ from incongruent features

We next compared the correlation between congruent and incongruent shape-sound features within features runs (*Figure 4a*). Complementing the previous behavioral pattern similarity results (*Supplemental Material Figure S1, S2*), we observed a main effect of learning day (before or after multimodal learning: *F*_1,32_ = 4.63, *p* = 0.039, *η*^2^ = 0.13), a main effect of congruency (congruent or incongruent object: *F*_1,64_ = 7.60, *p* = 0.0076, *η*^2^ = 0.11), and an interaction between learning day and congruency in the temporal pole (*F*_1,64_ = 6.09, *p* = 0.016, *η*^2^ = 0.087). Before multimodal learning, there was no difference in pattern similarity between congruent features compared to incongruent features (*t*_33_ = 0.22, p = 0.82). After multimodal learning, there was lower pattern similarity for shape and sound features associated with congruent compared to incongruent objects (*t*_33_ = 3.47,*p* = 0.0015, *Cohen’s d* = 0.22); *Figure 4*.

**Figure 4.**
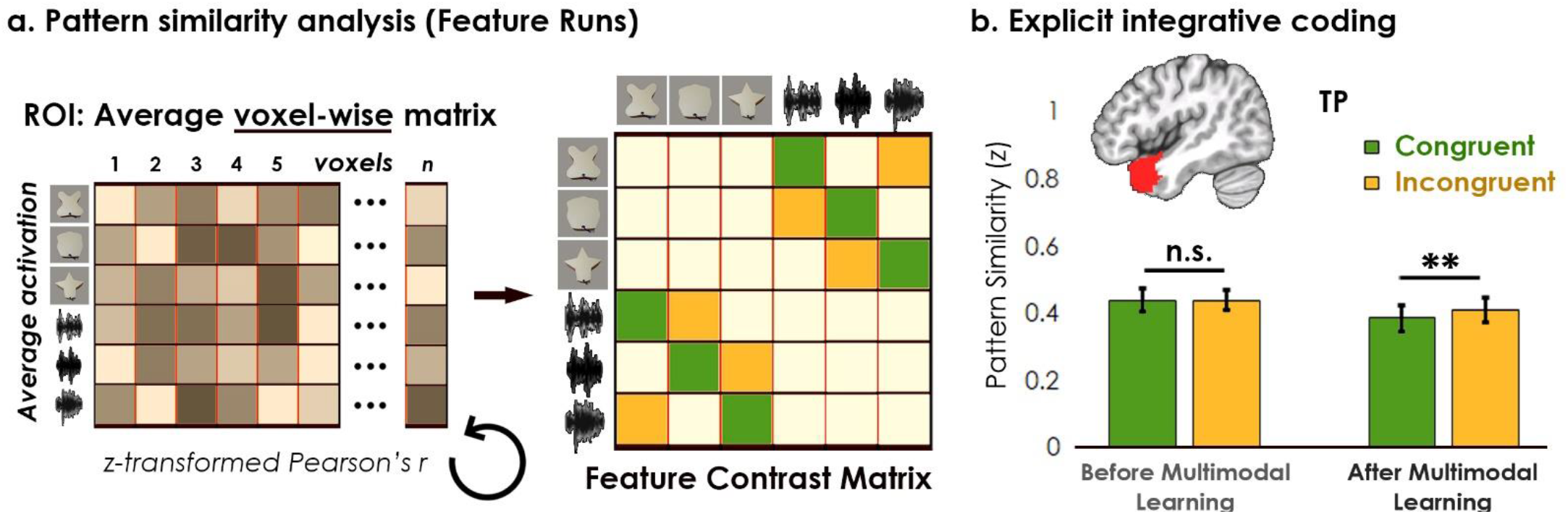
(*a*) Contrast matrix for feature runs shown on the left panel, and (*b*) actual pattern similarity analysis on the right. (*a*) The voxel-wise matrix averaged across feature runs was autocorrelated using the z-transformed Pearson’s correlation, creating a feature-level contrast matrix. We examined the average pattern similarity between features associated with congruent objects (green) compared to the same features associated with incongruent objects (yellow). (*b*) We found an interaction between learning day and congruency in the temporal pole (TP). Before multimodal learning, there was no difference in neural similarity between features associated with congruent objects compared to the same features associated with incongruent objects. After multimodal learning, there was *less* neural similarity between features associated with congruent objects compared to incongruent objects. Because congruent and incongruent objects are built from the same shapes and sounds, this result provides evidence of a multimodal object code that is different from the sum of the feature parts. ** denotes *p* < 0.01.

Although we observed greater pattern similarity between congruent compared to incongruent features in behavior (*Supplemental Material. Figure S2*), this *greater behavioral similarity* was related to *reduced neural similarity* following multimodal learning. These results provide neural evidence that features associated with congruent objects differ from the same features associated with incongruent objects in the temporal pole, providing evidence of an explicit integrative code (i.e., the whole object is different than the sum of the feature parts). By contrast, we did not observe an interaction between learning day and congruency in perirhinal cortex, LOC, V1, or A1 (*F*_1,60~64_ = 0.039~1.30, *p* = 0.26~0.84). Importantly, these pattern similarity analyses directly rule out a global novelty signal in anterior temporal lobe *per se*, as we observed robust neural changes differentiating congruent and incongruent objects even when participants experienced each feature sequentially during feature runs (*Figure 2, 4*).

#### The visually-biased code in perirhinal cortex was attenuated with learning

The previous analyses found evidence of an explicit integrative object-based code that was different from the object’s component features in the temporal pole. Next, we characterized how the individual feature representations may change when paired with different multimodal objects. One possibility is that the feature representations are not affected by object features from a different modality. For example, the visual representation of “frog” may remain unaffected by whether “frog” is paired with the sound of croaking or paired with the sound of barking (i.e., evidence that objects are constructed from sensory features). Alternatively, the feature representations may transform once paired with object features from a different modality. For example, the visual representation of “frog” may change when paired with the sound of croaking compared to barking (i.e., the whole object is abstracted away from component features).

First, the voxel-wise activity for *feature* runs was correlated to the voxel-wise activity for *object* runs before multimodal learning (*Figure 5a*). Consistent with previous univariate results (*Figure 3*), we observed greater pattern similarity for sound features in the temporal pole (*F*_1,32_ = 15.80,*p* < 0.001, *η*^2^ = 0.33) and A1 (*F*_1,32_ = 145.73, *p* < 0.001, *η*^2^ = 0.82), and greater pattern similarity for visual shape features in the perirhinal cortex (*F*_1,32_ = 10.99, *p* = 0.0023, *η*^2^ = 0.26), LOC (*F*_1,30_ = 20.09, *p* < 0.001, *η*^2^ = 0.40) and V1 (*F*_1,32_ = 22.02, *p* < 0.001, *η*^2^ = 0.41). Pattern similarity for each brain region was higher for one of the two modalities, indicative of a modality-specific bias towards either visual or sound content. By example, our results mean that the visual representation of “frog” was unaffected by the sound of croaking or by the sound of barking *before multimodal learning,* when a child has not yet learned the concept of frog.

**Figure 5.**
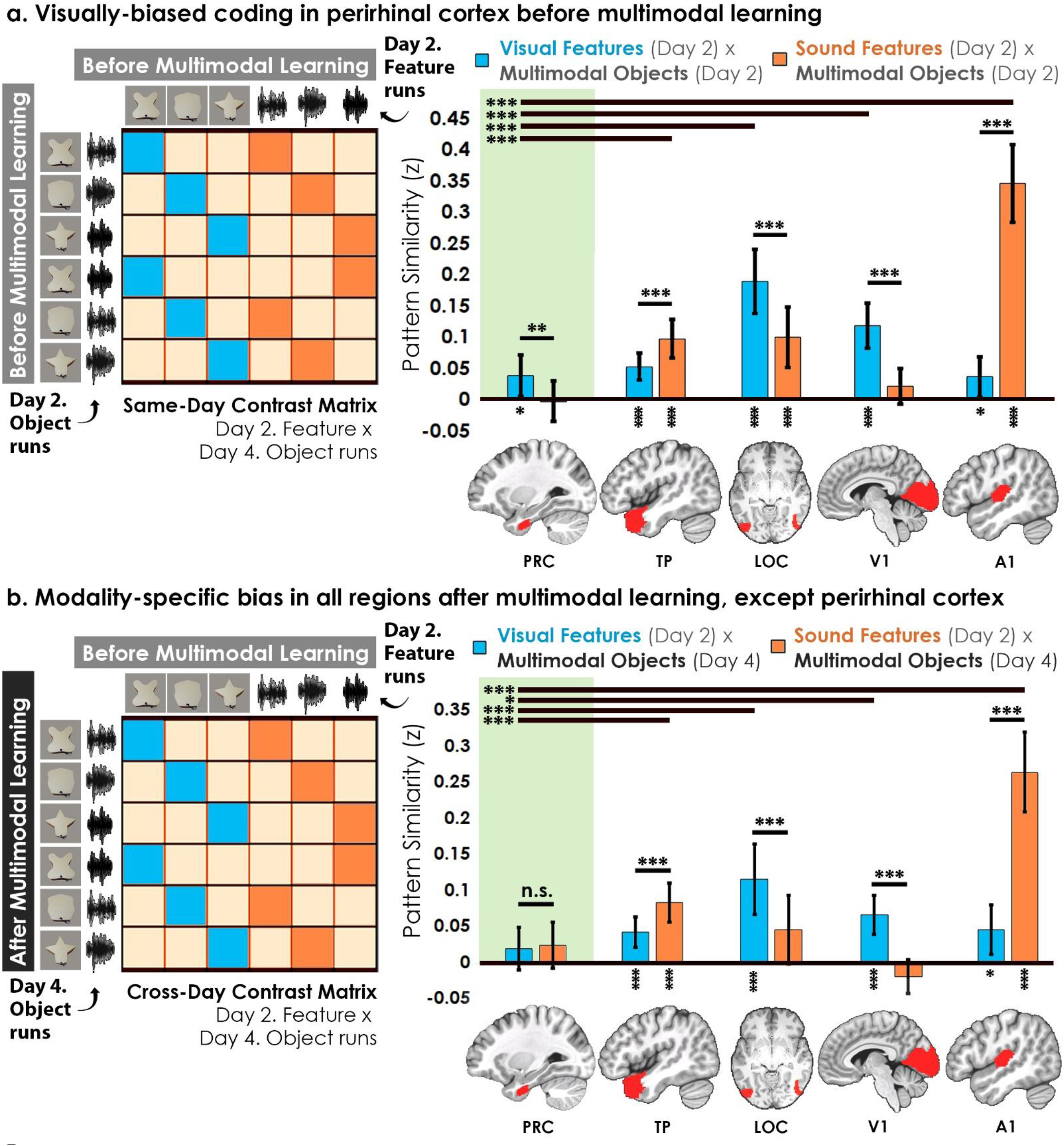
Contrast matrix shown on the left panels, with actual results shown on the right panels. The voxel-wise matrix for feature runs on Day 2 were correlated to the voxel-wise matrix for object runs on Day 2 (*a*) and Day 4 (*b*), creating a contrast matrix between visual and auditory features to different multimodal objects. We compared the average pattern similarity (z-transformed Pearson correlation) between shape (blue) and sound (orange) features across learning days. (*a*) Robust modality-specific feature biases were observed in all examined regions before multimodal learning. Pattern similarity for each brain region was higher for one of the two modalities, indicative of a modality-specific bias. For example, pattern similarity in perirhinal cortex (PRC) preferentially tracked the visual features of the multimodal objects, evidence of a default visual shape bias *before multimodal learning.* (*b*) Critically, we found that perirhinal representations were transformed with experience, such that the initial visual bias was attenuated *after multimodal learning* (i.e., denoted by a significant interaction, shown by shaded green regions), evidence that representations were no longer grounded in a single modality. ** denotes *p* < 0.05, ** denotes *p* < 0.01, *** denotes *p* < 0.001. Horizontal lines across brain regions indicate a significant effect of region relative to PRC, whereas horizontal lines within brain regions indicate a significant main effect of modality. Vertical asterisks denote pattern similarity comparisons relative to 0.

We then examined whether the original feature representations may change after participants learn that features are associated with specific multimodal objects (i.e., after multimodal learning, *Figure 2*). The voxel-wise activity for feature runs *before* multimodal learning was correlated to the voxel-wise activity for object runs *after* multimodal learning (*Figure 5b*). We found a significant interaction between modality and region (*F*_4,284_ = 27.64, *p* < 0.001, *η*^2^ = 0.28), meaning that the examined regions differed in their bias towards modality-specific features. Follow up contrasts revealed that the perirhinal cortex was no longer biased towards visual shape after multimodal learning (*F*_1,32_ = 0.12, *p* = 0.73), whereas the temporal pole, LOC, V1, and A1 remained biased towards either visual shape or sound (*F*_1,30-32_ = 16.20~73.42,*p* < 0.001, *η*^2^ = 0.35~0.70).

As the perirhinal cortex was the only region that differed in its modality-specific bias across learning days, we conducted a linear mixed model to directly track how perirhinal representations change from before multimodal learning to after multimodal learning (green shaded regions in *Figure 5a* and *5b*). This analysis revealed a significant interaction between learning day and modality (*F*_1,775_ = 5.56, *p* = 0.019, *η*^2^ = 0.071), meaning that the original visual shape bias observed in perirhinal cortex (*Figure 5a*) was attenuated with experience (*Figure 5b*). After multimodal learning, the same shape no longer invoked significant pattern similarity between objects that differed in terms of what they sounded like. By example, our results mean that *after* a child has learned that “frog” goes with croaking, there was no longer significant pattern similarity between “frog”-croaking with “frog”-barking – even though the same “frog” representation was carried over across feature pairings *before* multimodal learning. That is, we observed experience-dependent changes in perirhinal cortex as a function of multimodal learning, even though the physical features themselves had not changed. These results suggest that the perirhinal cortex changed how it tracked shape features with experience, evidence of an explicit integrative object representation transformed from the original features.

#### Experience-dependent explicit integrative coding in perirhinal cortex

So far, we have shown that the perirhinal cortex was by default biased towards visual shape features (*Figure 5a*), and that this visual shape bias was attenuated with experience (*Figure 5b*). Whereas the previous analysis tracked how features change when *paired with* other features (*Figure 5*), our final analyses tracked how the *individual object representations* themselves change after multimodal learning (*Figure 6*). Specifically, we compared the pattern similarity of objects across learning days, akin to examining how the combined “frog”-croaking representation itself might change after multimodal learning.

**Figure 6.**
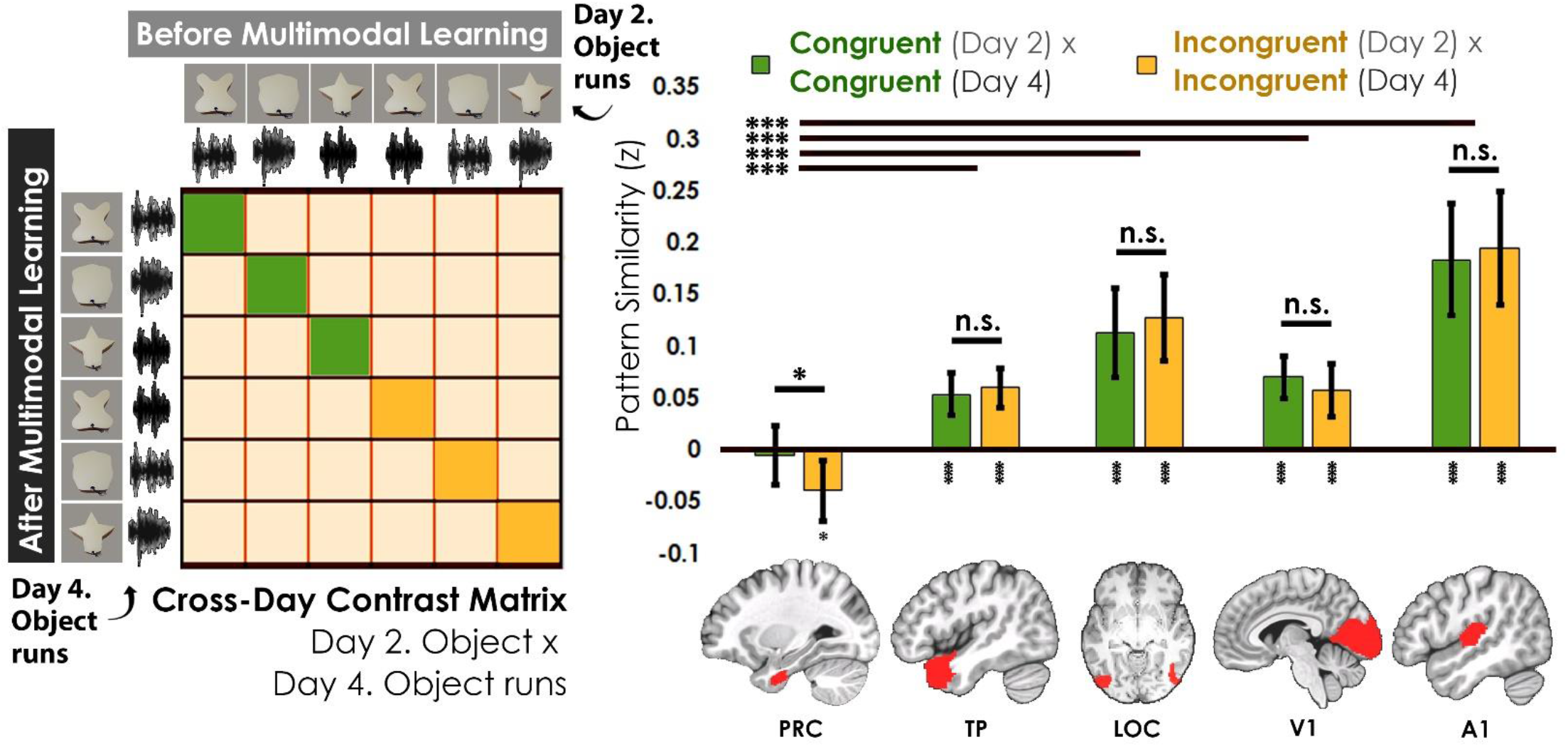
Contrast matrix shown on the left panel, with actual results shown on the right panel. We compared the average pattern similarity across learning days between object runs on Day 2 with object runs on Day 4 (z-transformed Pearson correlation). Critically, we observed lower average pattern similarity for incongruent objects (yellow) compared to congruent (green) objects in perirhinal cortex (PRC). These results suggest that perirhinal cortex differentiated congruent and incongruent objects constructed from the same features, with evidence of pattern separation that orthogonalized overlapping feature input into distinct object-level outputs (i.e., pattern similarity equal to or below 0). By contrast, there was no significant difference between congruent and incongruent objects in any other examined region. Furthermore, pattern similarity in TP, LOC, V1, and A1 were significantly above 0, suggesting that the representations from before multimodal learning were carried over after multimodal learning in these regions. * denotes *p* < 0.05, ** denotes *p* < 0.01, *** denotes *p* < 0.001. Horizontal lines spanning brain regions denote an effect of region relative to perirhinal cortex, whereas horizontal lines within brain regions denote a main effect of congruency. Vertical asterisks denote pattern similarity comparisons relative to 0.

Assessing the cross-day pattern similarity between object runs to object runs before and after learning (*Figure 6*), we observed a robust main effect of region (*F*_4,302_ = 47.71, *p* < 0.001, *η*^2^ = 0.39). Consistent with previous results (*Figure 3, 5*), perirhinal cortex representations differed from temporal pole, LOC, A1, and V1 (all *p* < 0.001). Furthermore, perirhinal cortex representations did not correlate significantly across learning days for objects (i.e., pattern similarity was never above 0), suggesting that the same “frog”-croaking representation from *before* multimodal learning transformed *after* multimodal learning.

Although pattern similarity was never above 0 across learning days, perirhinal cortex was the only region that differentiated between congruent and incongruent objects in this analysis (PRC: *F*_1,34_ = 4.67, *p* = 0.038, *η*^2^ = 0.12). Pattern similarity in perirhinal cortex did not differ from 0 for congruent objects across learning days (*t*_35_ = 0.39, *p* = 0.70) but was significantly lower than 0 for incongruent objects (*t*_35_ = 2.63, *p* = 0.013, *Cohen’s d* = 0.44). By contrast, pattern similarity in temporal pole, LOC, V1, and A1 was significantly correlated across learning days (pattern similarity > 0; all *p* < 0.001) and did not differ between congruent and incongruent objects (temporal pole, LOC, V1, and A1; *F*_1,32~34_ = 0.67~2.83, *p* = 0.10 ~ 0.42). In other words, perirhinal cortex not only differentiated between congruent and incongruent objects built from the same features (i.e., the whole object different from the sum of feature parts), but the original perirhinal representations changed with experience (i.e., denoted by the lack of significant pattern similarity above 0 for the same representations across learning days).

Taken together, these results critically suggest that multimodal learning transformed the original perirhinal representations (*Figure 5, 6*), forming coherent multimodal object representations that were abstracted away from the original component features.

### Discussion

Known as the *multimodal binding problem,* a long-standing question in the cognitive sciences has asked how the mind forms coherent multimodal concepts. To study this problem, we designed a 4-day task to decouple the learned multimodal object representations (Day 3 and 4) from the baseline unimodal shape and sound features (Day 1 and 2). Paired with multi-echo fMRI (*Figure 2*), our novel paradigm tracked the emergence of multimodal object concepts from component unimodal features in healthy adults. We found that the temporal pole and perirhinal cortex came to represent new object concepts with learning, such that the acquired object representations differed from the representation of the constituent features (*Figure 3, 4, 5, 6*). Intriguingly, the perirhinal cortex was by default biased towards visual shape, with this initial visual bias attenuated with experience (*Figure 3, 5*). Within the perirhinal cortex (*Figure 5, 6*), the acquired object concepts (measured after multimodal learning) were transformed away from original component features (measured before multimodal learning), providing evidence of experience-dependent changes following learning. These results reveal that sounds can be integrated with existing shape representations in the perirhinal cortex, suggesting that this region can serve as a polymodal integration hub for sensory input.

As one solution to the multimodal binding problem, we show that the temporal pole and perirhinal cortex form unique multimodal object representations distinct from features distributed across sensory regions (*Figure 3c, 4, 6*). In the temporal pole however, the multimodal object representations measured after multimodal learning were found to be related to the component feature representations measured before multimodal learning (*Figure 5, 6*). Thus, our results extend existing theoretical models,^2,3,4,5,7,8,9,27^ suggesting that the temporal pole may represent new multimodal objects by combining previously learned knowledge. For example, research into *conceptual combination* has linked the anterior temporal lobes to compound object concepts such as “hummingbird”.^28,29,30^ Participants during our task may have represented the sound-based “humming” concept and visually-based “bird” concept on Day 1, forming the multimodal “hummingbird” concept on Day 3; *Figure 1, 2.* For these reasons, the temporal pole may form a sparse multimodal object code based on pre-existing knowledge, resulting in reduced neural activity (*Figure 3e*) and pattern similarity after multimodal learning (*Figure 4b*).

By contrast, the perirhinal cortex represented the whole multimodal object distinct from the original component features. Whereas the temporal pole may combine pre-existing concepts (e.g., “hummingbird” from pre-existing “humming” and “bird” concepts), perirhinal cortex may orthogonalize overlapping feature input by creating distinct object representations through the operation of pattern separation.^5,6^ Indeed, there was no pattern similarity between the same congruent objects and *negative pattern similarity* between the same incongruent objects across learning days (*Figure 6*). On our task, participants must differentiate congruent and incongruent objects constructed from the same three shape and sound features (*Figure 2*). An efficient way to solve this task would be to form distinct object-level outputs from the overlapping feature-level inputs through a nonlinear transformation (i.e., measured as pattern similarity equal to or less than 0; *Figure 5b, 6*). Because our paradigm could decouple neural responses to the learned object representations (on Day 4) distinct from the original component features (on Day 2), these results provide evidence of pattern separation in the human perirhinal cortex.^5,6^

Importantly, we found that the anterior temporal lobe structures, including the perirhinal cortex, represented multimodal objects on the final day of our task after a period of multimodal learning (*Figure 2, 6b*). Although this observation is consistent with previous animal research^31^ finding that a period of experience is necessary for the perirhinal cortex to represent multimodal objects, future work will need to determine whether our findings are driven by *only* experience or by experience *combined with* sleep-dependent consolidation.^32^ Perhaps a future study could explore how separate features and the integrative object representations change over the course of the same learning day compared to multiple learning days after sleep. Nevertheless, perirhinal cortex was critically influenced by experience, potentially explaining why findings in this literature have been at times mixed, as stimulus history was not always controlled across different experiments.^33,34^ In our study, we explicitly controlled for stimulus history (*Figure 2*), ensuring that participants extensively explored individual features by the end of the first day and formed multimodal objects by the end of the third day.

Complementing seminal patient work causally linking anterior temporal lobe damage to the loss of object concepts,^35^ we show that the formation of new multimodal concepts also depends on the anterior temporal lobe. An important direction of future work may investigate the fine-grained functional divisions within the heterogeneous anterior temporal lobe region. One recent study has found that the anterior temporal lobe can be separated into 34 distinct functional regions,^36^ suggesting that a simple temporal pole versus perirhinal cortex division may not fully capture the complexity of this region. Imaging the anterior temporal lobe has long been known to be challenging with functional neuroimaging due to signal dropout.^37^ We show that a multi-echo fMRI sequence^20^ may be especially useful in future work, as multi-echo fMRI mitigates signal dropout better than the standard single-echo fMRI (see *Supplemental Material. Figure S3* for a comparison). Indeed, we observed robust anterior temporal lobe activity using a multi-echo sequence during a standard object functional localizer (*Figure 3c*), which typically finds activity only in earlier areas of the ventral visual stream using single-echo fMRI.^38^

Intriguingly, the initial visual shape bias observed in the perirhinal cortex was mitigated by experience (*Figure 5*). An important line of future investigation may be to explore whether the same experience-dependent changes occur in artificial neural networks that aim to learn multimodal object concepts.^39,40^ Previous human neuroimaging has shown that the anterior temporal lobes are important for intra-object configural representations,^41^ such that damage to the perirhinal cortex^14,42^ leads to object discrimination impairment. For example, human participants with perirhinal cortex damage are unable to resolve feature-level interference created by viewing multiple objects with overlapping features. Certain types of errors made by deep learning models^43^ also seem to resemble the kinds of errors made by human patients,^14,35,44^ whereby accurate object recognition can be disrupted by feature-level interference. Writing the word “iPod” on an apple image, for instance, can lead to deep learning models falsely recognizing the apple as an actual iPod.^45^ Existing artificial neural network models also do not yet reach human performance on tasks involving multimodal integration.^39,40^ For these reasons, future work to mimic the coding properties of anterior temporal lobe structures in artificial neural networks may enable machines to better approximate the remarkable human ability to learn concepts.

In summary, forming multimodal object concepts relies on the representations for the whole object in anterior temporal lobe structures distinct from the distributed feature representations in sensory regions. It is this hierarchical architecture that supports our ability to understand the external world, providing one solution to the age-old question of how multimodal concepts can be constructed from their component features.

## Acknowledgements

We are grateful to the Toronto Neuroimaging community for helpful feedback. In particular, the first author thanks Dr. Katherine Duncan and Dr. Massieh Moayedi for suggestions related to the experimental design, Dr. Michael Mack for initial guidance with 3D-printing, Dr. Rosanna Olsen for her tutorial on medial temporal lobe segmentation, as well as Dr. Andy Lee and Dr. Adrian Nestor for their neuroimaging and multivariate pattern analysis courses.

We thank Annie Kim and Katarina Savel for their assistance with participant recruitment, as well as Priya Abraham for her assistance with MRI scanning. Finally, we thank Dr. Rüdiger Stirnberg for sharing with us the multi-echo fMRI sequence used in this manuscript.

AYL is supported by an Alexander Graham Bell Canada Graduate Scholarship-Doctoral from the Natural Sciences and Engineering Research Council of Canada (NSERC CGS-D). This work is supported by a Scholar Award from the James S McDonnell Foundation, an Early Researcher Award from the Ontario Government, an NSERC Discovery grant, and a Canada Research Chair to MDB.

## Methods

All experiments described in this study were approved by the University of Toronto Ethics Review Board: 37590.

### Stimulus Validation Experiment

#### Participants

16 participants were recruited from the University of Toronto undergraduate participant pool and from the community. Course credit or $10/hr CAD was provided as compensation.

#### Validation Procedure

The stimulus validation procedure was based on previous work.^21^ Participants rated the similarity of each stimulus in the context of each other stimulus 8 times, with 4 repeats of the same stimulus. In a self-timed manner, participants viewed pictures of shapes or clicked icons to play the to-be-rated sounds from a headset. The specific shapes were sampled from a triangular geometry on the *Validated Circular Shape Space,*^21^ and we replicated the subsequent triangular geometry from participant similarity ratings (*Figure 1a;* each stimulus was about as similar as every other stimulus). On a separate task, the same participants rated the subjective similarity between five experimenter-created sounds, and we selected three which formed a triangular geometry. This procedure defined the *a priori* representational geometry of the individual shapes and sounds, ensuring that subjective similarity was well-matched prior to the learning task.

#### 3D-Printing

The three validated shapes were 3D-printed using a DREMEL Digilab 3D Printer 3D45-01 with 1.75 mm gold-colored polymerized lactic acid filament. To create the 3D object models, the original 2D images were imported into Blender and elongated to add depth. The face of the shape image created a detachable lid, with a small circular opening included to allow wiring to extend to a playable button positioned on the exterior of the shape. An empty space was formed inside the 3D shape for the battery-powered embedded speaker. To ensure that the objects were graspable, each shape was 3D-printed to be approximately the size of an adult hand (*Figure 1c*). The lid of the shape was detached before each learning day (*Figure 2*), with the embedded speaker programmed to play either no sound or a paired sound from the validated set of stimuli (*Figure 1a*). After the speaker was programmed, the lid of the shape was reattached using thermoplastic adhesive.

### 4-Day Multimodal Object Learning Task using Multi-Echo fMRI

#### Participants

Compensation was $250 CAD for the 2 neuroimaging sessions and 2 behavioral sessions (~approx. 6 hours total, which included set-up, consent, and debriefing), with a $50 CAD completion bonus. 20 new participants were recruited and scanned at the Toronto Neuroimaging Facility.

All participants were right-handed, with normal or corrected-to-normal vision, normal hearing, and no history of psychiatric illness. Of the 20 scanned participants, 1 participant dropped out after the first neuroimaging session. Severe distortion was observed in a second participant from a metal retainer and data from this participant was excluded from subsequent analyses. Due to technical difficulties, the functional localizer scans were not saved for one participant and most feature runs could not be completed for a second participant. Overall, the within-subject analyses described in the main text included data from at least 16 participants, with most analyses containing data from 17 participants across two scanning days (each scanning session lasting approximately 90 minutes).

#### Behavioral Task

On each behavioral day (*Figure 2*), participants explored the stimuli and then provided similarity judgments on a separate task. As practice for the neuroimaging days, participants also completed one feature-and one object n-back task after the initial exploration phase.

#### Exploration Phase

On Day 1 (*Figure 2a*), participants separately learned the shape and sound features in a random order. The 3D shapes were explored and physically palpated by the participants. We also encouraged participants to press the button on each shape, although the button was not operational on this day. Each 3D-printed shape was physically explored for 1 minute and each sound was heard through a headset 7 times. The exploration phase was repeated after each task and after every 24 trials of the similarity rating task. This procedure ensured that each individual stimulus was experienced extensively by the end of the first day.

On Day 3 (*Figure 2c*), participants experienced the shape and sound features together by pressing the now-operational button on each object, with multimodal objects explored in a random order. Participants were allotted 1 minute to physically explore and palpate each multimodal object, as well as to listen to the associated sound by pressing the button. After the initial exploration phase, participants completed practice feature and object *n*-back tasks, then followed by the similarity rating task. Identical to Day 1, the exploration phase was repeated between tasks and after every 24 trials of the pairwise similarity rating task.

#### Pairwise Similarity Task

Using the same task as the stimulus validation procedure, participants provided similarity ratings for all combinations of the 3 shapes and 3 sounds (rated 8 times in the context of each other stimulus in the set, with 4 repeats of the same item; 72 total trials).

#### Multi-echo fMRI Data Acquisition

A 3D multi-echo EPI sequence with blipped-CAIPI sampling^46^ was used to acquire fMRI data on Day 2 and Day 4. For task-related scans, the 3 echoes (TR = 2000 ms, TE 1 = 11 ms, TE 2 = 31.6 ms, and TE 3 = 52.2 ms) were each acquired with 90 images (210 × 210 field of view with a 100 × 100 matrix resize; anterior to posterior phase encoding, 78 slices, slice thickness: 2.10 mm, flip angle: 17°, interleaved multi-slice acquisition), resulting in an in-plane resolution of 2.10 × 2.10 mm. 3D distortion correction and pre-scan normalization was enabled, with acceleration factor PE = 2 and acceleration factor 3D = 3. These parameters yielded coverage over the entire cortex, and a B0 field map was collected at the completion of the experiment.

A standard whole-brain high-resolution T1-weighted structural image was collected (TR = 2000 ms, TE = 2.40 ms, flip angle = 9°, field of view = 256 mm, 160 slices, slice thickness = 1.00 mm, acceleration factor PE = 2), resulting in an in-place resolution of 1.00 mm × 1.00. Two 6 minute 42 second resting state scans were also collected (TR = 2000 ms, TE = 30 ms; field of view: 220 mm, slice thickness: 2.00 mm; interleaved multi-slice acquisition, with acceleration factor PE = 2).

Scanning was conducted using a 32-channel receiver head coil with the Siemens Magnetom Prisma 3T MRI scanner at the Toronto Neuroimaging Facility. To record responses, participants used a 4-button keypad (Current Designs, HHSC-1X4-CR). Stimuli materials were displayed using an MR compatible screen at high resolution (1920 × 1080) with zero-delay timing (32” BOLD screen) controlled by PsychToolbox-3 in MATLAB. At the start of each neuroimaging session, we performed a sound check with a set of modified in-ear MR-compatible headphones (Sensimetrics, model S14), followed by a functional localizer and then by the task-related runs.

#### fMRI Experimental Procedure

Rather than collecting data from many different instances of a category as is common in a fMRI study using multivariate pattern analysis, we collected data from many repetitions of the *same* stimulus using a psychophysics-inspired approach. This paradigm ensured that the neural representations specific to each unimodal feature and each multimodal object was well-powered for subsequent pattern similarity analyses.^47^ Excluding *n*-back repeats, each feature was displayed 4 times per run for a total of 40 instances per scanning session (80 instances of each unimodal feature in total). Excluding *n*-back repeats, each shape-sound pairing was displayed 4 times per run for a total of 20 instances per scanning session (40 instances of each shape-sound object in total).

The primary task was divided across 10 feature runs and 5 object runs of functional data acquisition. Two feature runs followed by one object run were presented in an interleaved manner to participants until all 10 feature runs and 5 object runs were completed. After an initial functional localizer, we collected a resting state scan. After five task-related runs, we acquired a whole-brain high-resolution T1-weighted structural image. After 10 task-related runs, we acquired a second resting-state scan.

We designed our task-related runs to be 3 minutes in length, as “mini-runs” have been shown to improve data quality in multivariate pattern analysis.^47^ To begin each feature run, a blank screen appeared for 6 seconds. A shape or sound feature was then presented for two seconds, followed by a fixation cross appearing for 2 – 8 seconds (sampled from the following probability distribution: 2 seconds = 30%, 4 seconds = 30%, 6 seconds = 30%, and 8 seconds = 10%). Each stimulus was presented four times in a random order per run, with two repeats occurring at a random position for the *n*-back task. The stimulus identity and temporal position of any given *n-*back repeat was random.

For the object runs, each shape appeared on the monitor at the same time as a sound was played through the headset for two seconds. Ensuring equal trial numbers, three shape-sound pairings were congruent (learned by participants) and three shape-sound pairings were incongruent (not learned by participants). Congruent and incongruent pairings were built from different combinations of the same shape and sound features, with pairings counterbalanced across participants. The procedure for the object runs were the same as the feature runs, except participants were asked to respond to an exact sequential repeat of *both* shape and sound.

#### Standard Object Functional Localizer

Participants viewed intact objects and phase scrambled versions of the same objects in separate 24 second blocks (8 functional volumes).^38^ Each of the 32 images within a block were presented for 400 ms each with a 350 ms ISI. There were 2 groups of 4 blocks, with each group separated by a 12 s fixation cross. Block order was counterbalanced across participants. All stimuli were presented in the context of an *n*-back task, and the order of images within blocks was randomized with the 1-back repeat occurring once per block. The identity and temporal position of the *n*-back repeat was random.

#### ROI Definitions

We conducted region-of-interest univariate (*Figure 3d, e*) and multivariate pattern analysis (*Figure 4, 5, 6*) in five masks: temporal pole, perirhinal cortex, lateral occipital complex (LOC), primary visual cortex (V1), and primary auditory cortex (A1). Temporal pole, V1, and A1 masks were extracted from the Harvard-Oxford atlas. The perirhinal cortex mask was created from the average of 55 manually-segmented T1 images from a previous publication.^48^ The LOC mask was extracted from the top 500 voxels in the lateral occipital region of each hemisphere that activated more strongly to intact than phase scrambled objects in the functional localizer (uncorrected voxel-wise p < 0.001).^38^ Probabilistic masks were thresholded at .5 (i.e., voxels labelled in 50% of participants), with the masks transformed to subject space through the inverse warp matrix generated from FNIRT nonlinear registration (see *Preprocessing*) then resampled from 1mm^3^ to 2.1mm^3^. All subsequent analyses were conducted in subject space.

#### Multi-echo ICA-based Denoising

For a detailed description of the overall ME-ICA pipeline, see the *tedana* Community.^49^ The multi-echo ICA-based denoising approach was implemented using the function *meica.py* in AFNI.

We optimally combined the signal across echoes 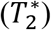 using a weighted average which accounts for higher signal in earlier echoes (i.e., lower TE) and higher sensitivity at later echoes (i.e., higher TE). PCA then reduced the dimensionality of the optimally-combined dataset and ICA decomposition was applied to remove non-BOLD noise. TE-dependent components reflecting BOLD-like signal for each run were used as the dataset for subsequent preprocessing in FSL.

#### Preprocessing

First, the anatomical image was skull-stripped. Data were high-pass temporally filtered (50 s) and spatially smoothed (6 mm). Functional runs were registered to each participant’s high-resolution MPRAGE image using FLIRT boundary-based registration, with registration further refined using FNIRT nonlinear registration. The resulting data were analyzed using first-level FEAT Version 6.00 in each participant’s native anatomical space.

#### Univariate Analysis

To obtain participant-level contrasts, we averaged the run-level feature (*Visual* > *Sound*) and object (*Congruent* > *Incongruent*) runs to produce the whole-brain group-level contrasts in FSL FLAME. Following standard recommendations, whole-brain analyses were thresholded at voxel-level *p* = 0.001 with random field theory cluster correction at *p* = 0.05.

For ROI-based analyses (*Figure 3d, e*), we estimated percent signal change using *featquery.* The parameter estimates (beta weight) were scaled by the peak height of the regressor, divided by the baseline intensity in the *Visual* > *Sound* and *Congruent* > *Incongruent* contrasts to obtain a difference score. Inferential statistical analyses were performed using a linear mixed model with learning day (before or after multimodal learning) and hemisphere (left or right) as fixed effects, with participants modelled as random effects. All subsequent linear mixed model analyses were conducted using the *nlme* package in R version 3.6.1.

#### Single Trial Estimates

We used the least squares single approach^50^ with 2 mm smoothing on the raw data in a separate set of analyses distinct from the univariate contrasts. Each individual stimulus, all other repetitions of the stimulus, and all other individual stimuli were modelled as covariates, allowing us to estimate whole-brain single-trial betas for each trial by run by mask by hemisphere by subject. All pattern similarity analyses described in the main text were conducted using the *CoSMoMVPA* package in MATLAB. After the single-trial betas were estimated, the voxel-wise activity across runs were averaged into a single overall matrix.

#### Behavioral Pattern Similarity Analysis

The behavioral ratings for each stimulus was averaged into a single feature-level RDM. We examined the magnitude of pattern similarity for congruent features compared to incongruent features across learning days.

#### Neuroimaging Pattern Similarity Analysis

Four comparisons were conducted: 1) the autocorrelation of the feature-level RDM (*Figure 4a*), 2) the correlation between the feature run RDM before multimodal learning to the object run RDM before multimodal learning (*Figure 5a*), 3) the correlation between the feature run RDM before multimodal learning to the object run RDM after multimodal learning (*Figure 5b*), and 4) the correlation between the object run RDM before multimodal learning to the object run RDM after multimodal learning (*Figure 6*).

The z-transformed Pearson’s correlation coefficient was used as the distance metric for all pattern similarity analyses. Inferential statistical analyses were performed using linear mixed models with congruency (congruent or incongruent), learning day (before or after multimodal learning), brain region (perirhinal cortex, temporal pole, LOC, V1, A1), modality (visual or sound), and hemisphere (left or right) as fixed factors, with participant modelled as random effects allowing intercepts to vary by learning day when appropriate. One-sample t-tests also compared the z-transformed pattern similarity scores relative to 0.

### Behavior-only Learning Task

#### Participants

44 new participants were recruited from the University of Toronto undergraduate participant pool and from the community. Course credit or $10/hr CAD was provided as compensation.

#### Procedure

In a same-day variant of the four-day task described in the main text (*Figure 2*), participants first explored the shapes and sounds separately. On a separate task, participants provided similarity ratings for all combinations of the 3 shapes and 3 sounds (rated 8 times in the context of each other stimulus in the set, with 4 repeats of the same item; 72 total trials). Every 24 trials, participants again explored the same shapes and sounds (separately before multimodal learning, in a counterbalanced order across participants). The behavioral similarity judgments were analyzed in the same pattern similarity approach described in the main text (*Supplemental Material. Figure S1, S2*).

