## Supplemental Material for "Multimodal Object Representations Rely on Integrative Coding"

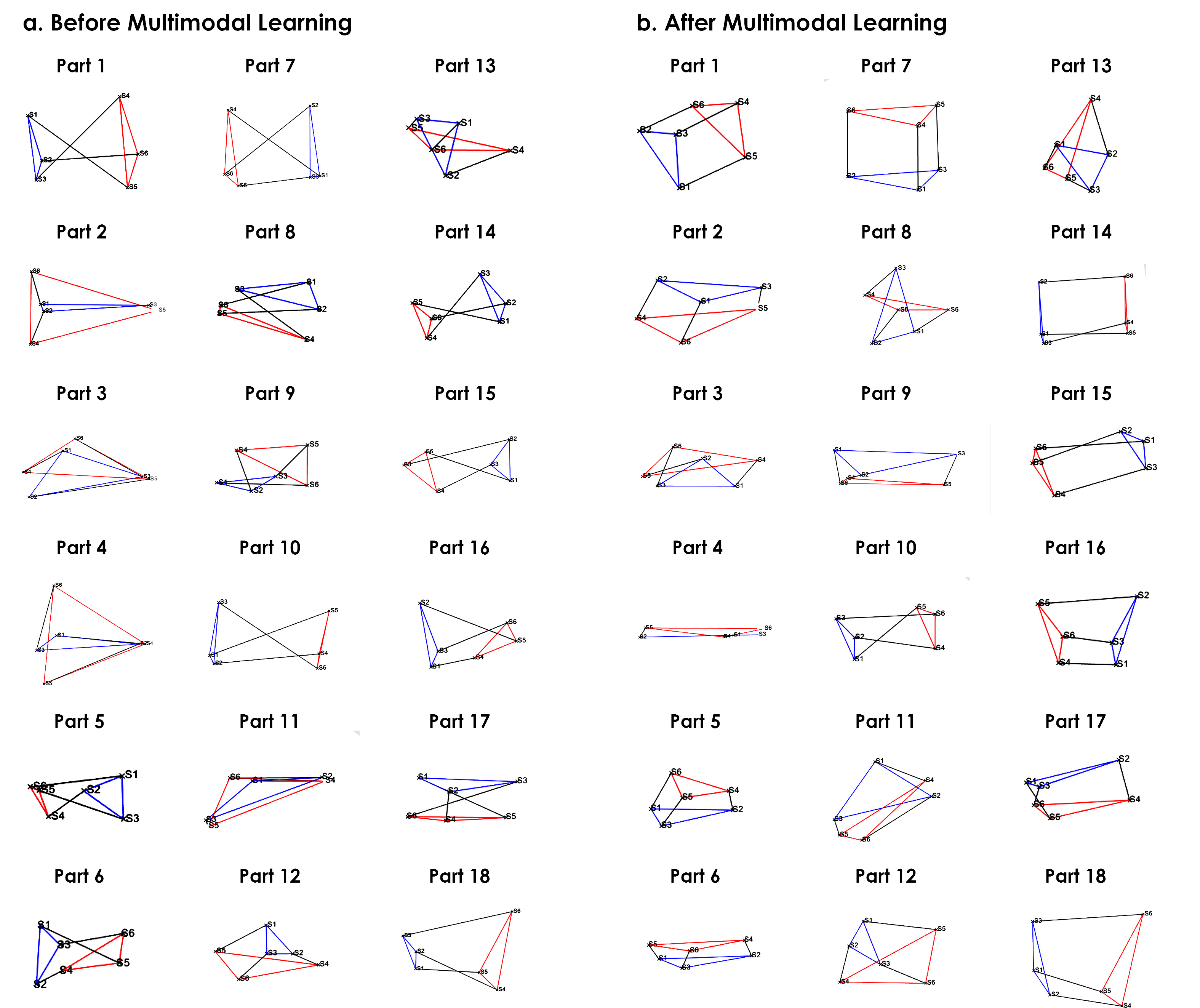

*Figure S1.* Participant representational geometries visualized from behavioral similarity ratings using multidimensional scaling. (*a*) Before multimodal learning, we replicated the triangular representational geometry of the shapes (blue) and sounds (red) sampled from the independent validation experiment. The congruent feature association to-be-learned on Day 3 is visualized as a line (black) connecting individual shape (blue) and sound (red) features. (*b*) After multimodal learning, participant representational geometries were now additionally driven by new information about the learned shape-sound associations. The shapes and sounds associated with congruent objects became more similar (i.e., closer together on the figure above) than the same shapes and sounds associated with incongruent objects, forming representational geometries that resembled a triangular prism after multimodal learning. These results indicate that multimodal learning changed how participants represented the shape-sound features, because participants experienced the same unimodal shapes and sounds across all four days of the experiment. That is, new information about the learned associations were now present in the representational geometry, evidence of an integrative code specific to the learned object (i.e., new information about the object measured after multimodal learning) distinct from the original feature components (i.e., the baseline representational geometries measured before multimodal learning).

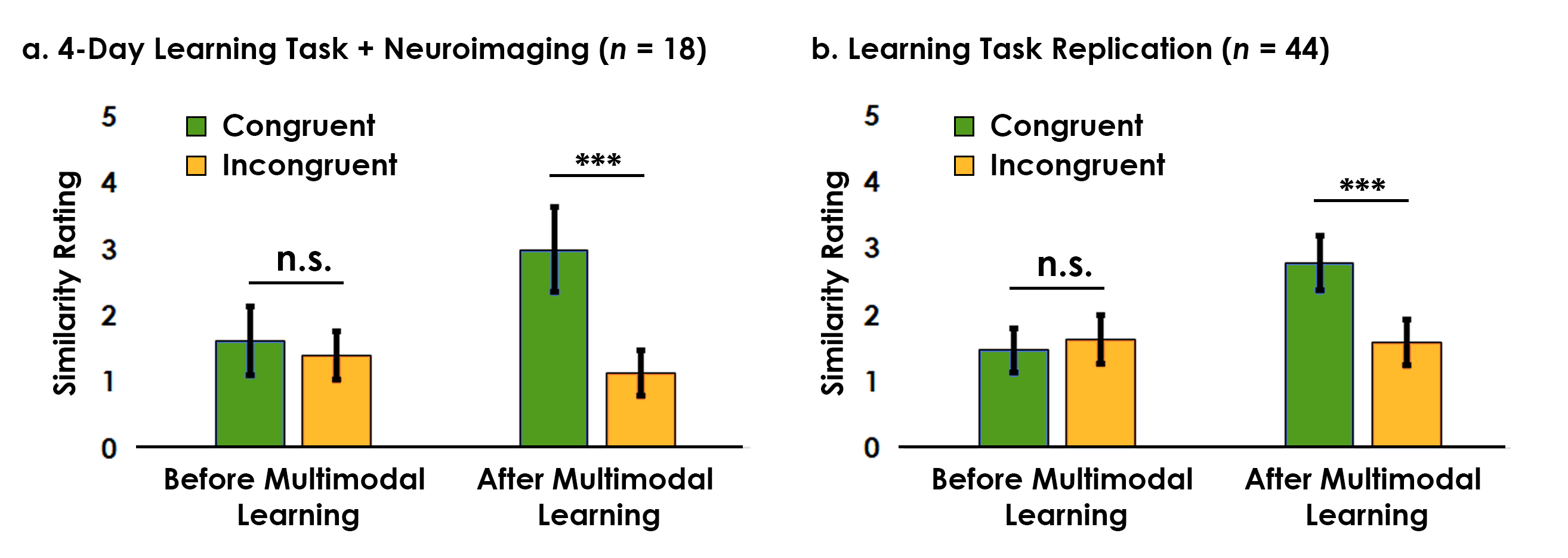

*Figure S2.* Behavioral pattern similarity results. (*a*) Results from the four-day learning task using multi-echo fMRI described in the main text. Before multimodal learning, there was no difference in similarity between shape and sound features associated with congruent objects compared to incongruent objects. After multimodal learning, we observed a robust shift in the magnitude of similarity. The shape and sound features associated with congruent objects were now significantly more similar than the same shape and sound features associated with incongruent objects (p < 0.001), evidence that multimodal learning changed how participants experienced the unimodal features (observed in 17/18 participants). (*b*) We replicated this learning-related shift in pattern similarity with a larger sample size (*n* = 44; observed in 38/44 participants). Across two experiments, we observed a robust experience-dependent shift in pattern similarity across 55/62 participants (88%). As the physical shape and sound features did not change across the learning days, the observed change in similarity provides robust evidence that multimodal learning influenced how participants subjectively experienced the shape-sound features.

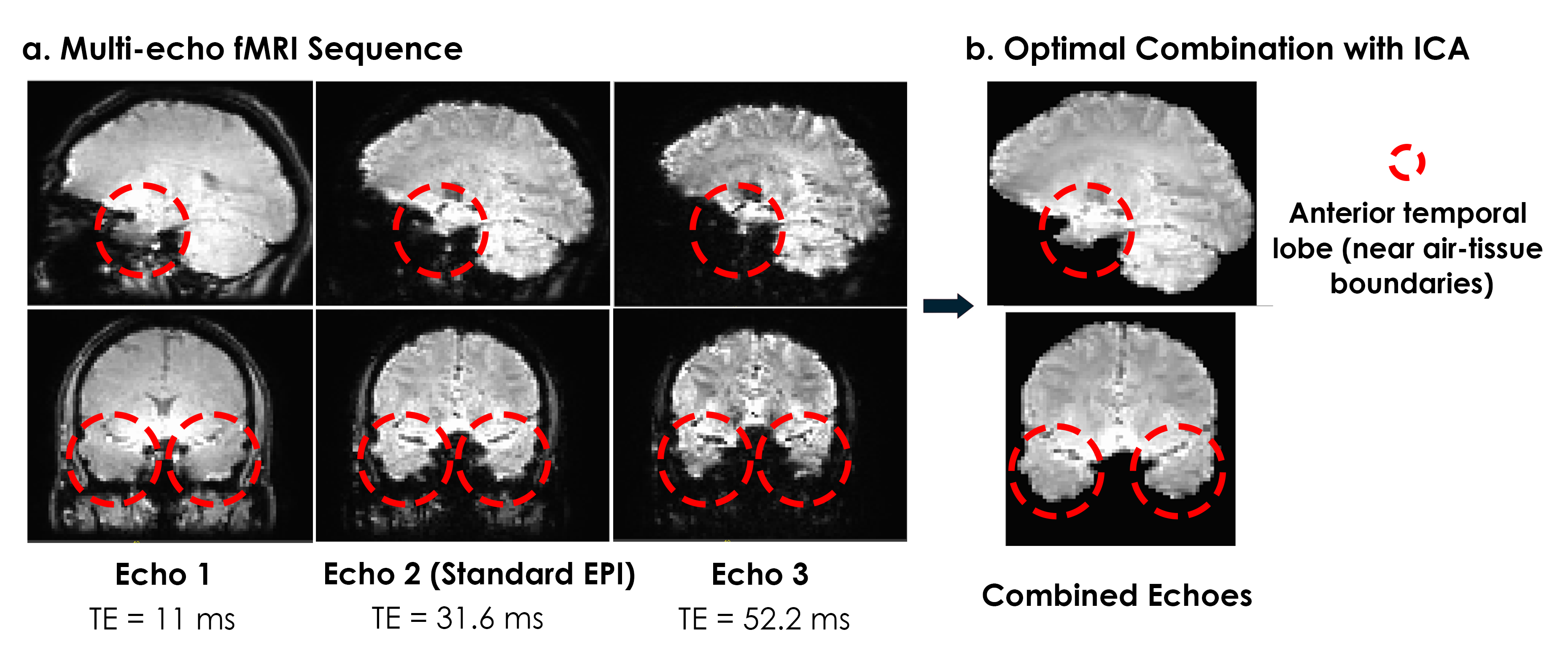

*Figure S3*. Signal quality comparison from a representative participant. (*a*) The multi-echo sequence we used acquired 3 measurements after every radiofrequency pulse, compared to the standard single-echo EPI which acquires a single measurement (usually at a TE around 30 ms). A multi-echo sequence with 3 echoes acquires 3 times as much data as the current standard single-echo approach, and accounts for differences in measured T2* across brain regions. For example, better signal is obtained at high TE values for the anterior temporal lobes, which would otherwise reveal substantial signal dropout due to susceptibility artifacts at TE = 30 ms. (*b*) We optimally combined the three echoes, and then applied automatic ICA-based denoising to remove non-BOLD related noise. We found that the multi-echo approach better recovers signal from the anterior temporal lobe structures compared to the standard single-echo approach, likely accounting for the robust anterior and medial temporal lobe activity observed in a standard functional localizer comparing intact to scrambled objects described in the main text (*Figure 3c*).
